# Identification of antibiotic resistance proteins via MiCId’s augmented workflow. A mass spectrometry-based proteomics approach

**DOI:** 10.1101/2021.11.17.468978

**Authors:** Gelio Alves, Aleksey Ogurtsov, Roger Karlsson, Daniel Jaén-Luchoro, Beatriz Piñeiro-Iglesias, Francisco Salvà-Serra, Björn Andersson, Edward R.B. Moore, Yi-Kuo Yu

**Affiliations:** National Center for Biotechnology Information, National Library of Medicine, National Institutes of Health, Bethesda, MD 20894, USA; Department of Infectious Diseases, Sahlgrenska Academy, University of Gothenburg, Gothenburg, Sweden; Department of Clinical Microbiology, Sahlgrenska University Hospital, Gothenburg, Sweden; Center for Antibiotic Resistance Research (CARe), University of Gothenburg, Gothenburg, Sweden; Nanoxis Consulting AB, Gothenburg, Sweden; Culture Collection University of Gothenburg (CCUG), Sahlgrenska Academy of the University of Gothenburg, Gothenburg, Sweden; Microbiology, Department of Biology, University of the Balearic Islands, Palma de Mallorca, Spain; Bioinformatics Core Facility at Sahlgrenska Academy, University of Gothenburg, Gothenburg, Sweden

**Keywords:** Identification of antibiotic resistance proteins, Microorganism identification/classification workflow, Mass spectrometry

## Abstract

Fast and accurate identifications of pathogenic bacteria along with their associated antibiotic resistance proteins are of paramount importance for patient treatments and public health. While mass spectrometry has become an important, technique for diagnostics of infectious disease, there is a need for mass spectrometry workflows offering this capability. To meet this need, we have augmented the previously published <u>Mi</u>croorganism <u>C</u>lassification and <u>Id</u>entification (MiCId) workflow for this capability. To evaluate the performance of the newly augmented MiCId workflow, we have used MS/MS datafiles from samples of 10 antibiotic resistance bacterial strains belonging to three different species: *Escherichia coli, Klebsiella pneumoniae*, and *Pseudomonas aeruginosa*. The evaluation results show that MiCId’s workflow has a sensitivity value around 85% (with a lower bound at about 72%) and a precision greater than 95% in the identification of antibiotic resistance proteins. Using MS/MS datasets from samples of two bacterial clonal isolates, one being antibiotic-sensitive while the other (obtained from the same patient at different times) being multidrug-resistant, we applied MiCId’s workflow to investigate possible mechanisms of antibiotic resistance in these pathogenic bacteria; the results showed that MiCId’s conclusions are in agreement with the published study. Furthermore, we show that MiCId’s workflow is fast. It provides microorganismal identifications, protein identifications, sample biomass estimates, and antibiotic resistance protein identifications in 6–17 minutes per MS/MS sample using computing resources that are available in most desktop and laptop computers, making it a highly portable workflow. This study demonstrated that MiCId’s workflow is fast, portable, and with high sensitivity and high precision, making it a valuable tool for rapid identifications of bacteria as well as detection of their antibiotic resistance proteins. The new version of MiCId (v.07.01.2021) is freely available for download at https://www.ncbi.nlm.nih.gov/CBBresearch/Yu/downloads.html.

## 1 Introduction

Fast and accurate identification of pathogenic bacteria along with the identification of antibiotic resistance (AR) proteins is of paramount importance for patient treatments and public health [1–5]. Once the pathogenic bacteria causing the infections are identified swiftly along with their AR proteins (if present), proper treatment can be administered which can increase patients’ survival rate and minimize improper use of antibiotics [6,7].

Currently, molecular methods such as next-generation sequencing (NGS) and mass spectrometry (MS) are used and are being developed to speed up identifications of pathogenic bacteria [8–25]. While several computational workflows/pipelines for analyzing NGS data have been developed to identify pathogenic bacteria and AR genes [26–28], a mass spectrometry workflow with this capability is still lacking [24]. This has motivated us to augment the workflow of our pathogen identification tool, <u>Mi</u>croorganism <u>C</u>lassification and <u>Id</u>entification (MiCId) [21,29,30], to enable the identification of AR proteins, using MS data from a high-performance liquid chromatography system coupled to a high resolution tandem MS (HPLC-MS/MS). Another motivation for augmenting MiCId’s workflow is that, even though NGS workflows can provide information about the presence of AR genes, they do not provide information about protein expression, which is extremely important for treating infections and for understanding the mechanism of antibiotic resistance in bacteria [31–34].

For a summary of some of the existing workflows employed for the identification of bacteria using HPLC-MS/MS experiments, we refer readers to previous publications [24,29,30]. Overall, there has been significant progress made in the identification of bacteria using HPLC-MS/MS experiments, although there is plenty of room for improvement in sample preparation protocols and data analysis workflows [35–37]. Developers of HPLC-MS/MS data analysis workflows often use the sensitivity (true positive rate) and specificity (true negative rate) as the only criteria to assess the usability of the developed workflow. Although sensitivity and specificity are acceptable criteria to measure the performance of a workflow, these criteria alone are not enough to justify the usability of a workflow. For example, an important criterion that is often not mentioned in performance evaluations is the execution time. Identification of bacteria is a computationally demanding task for a workflow, as it has to query tens of thousands of MS/MS spectra in a microorganismal database containing thousands to tens of thousands of bacteria. In order to scale with the number of HPLC-MS/MS experiments, a workflow with appropriate amount of computer resources must have execution time less than the time it takes to conduct the HPLC-MS/MS experiment, which is approximately 1-2 hours. This remained an unattainable goal for most workflows [38]. Other criteria to consider include whether or not a workflow provides for identified bacterial biomass estimation [39,30], protein identification [40,29] with protein quantification [41,42], and AR protein identification [24]. Data on the relative biomasses of identified bacteria identified are essential for studying microbial communities [39] and are valuable when determining treatment options for patients suffering from co-infections [43–45]. Knowledge of proteins and protein expression levels are essential for analyzing gene expression and function [46,47] and for investigating possible mechanisms of antibiotic resistance in bacteria [31–34]. Information about AR proteins is crucial for proper treatments for AR-resistant bacterial infections [6,7]. The criteria above cover most of the data analysis features needed for a workflow to be useful. In order to ensure a workflow to be user-friendly, intuitive, and customizable, we propose additional criteria. A useful workflow should: (1) automate and customize microorganismal protein sequences for download and database construction; (2) automate and customize AR protein sequences for download and database construction; (3) be computationally efficient and scalable to handle large microorganismal databases, large numbers of MS/MS spectra, and large number of MS/MS experiments; (4) be available to execute in different computer operating systems; (5) offer a user-friendly graphical interface. Meeting these latter criteria allows a workflow to eliminate elaborate intermediate steps and reaching a broader group of users in addition to experts in the field.

In previous studies, we have demonstrated that MiCId’s workflow meets most of the criteria listed above [21,29,30,48]. We have shown that MiCId’s workflow:

Offers automated microorganismal database construction by automatically downloading from the NCBI database protein sequences of organisms specified by the user.
Offers customized microorganismal database construction, using a list of protein sequence Fasta files of organisms specified by the user that are stored in the local computer.
Is able to identify bacteria in samples containing single and multiple bacteria with high sensitivity and high specificity by computing, for each identified taxon, an *E*-value which can be used to control the proportion of false discoveries (PFD) without the need of a decoy database [21,29]. When a list of candidate taxa are ranked by a quality score *S*, the *E*-value *E*(*S* ≥ *S*_0_) should is defined as the expected number of random taxa with scores the same as or better than *S*_0_.
Is able to estimate taxonomic biomass by computing a quantity called the *prior* using a modified expectation-maximization (EM) method. The *prior* is defined as the probability for a taxon to emit any evidence peptide and can be regarded as the taxon’s relative protein biomass within the sample analyzed [30].
Provides protein identifications via combining peptides’ *E*-values, using theoretically derived mathematical formulas [49,40].
Is computationally efficient and scalable, taking 6-17 minutes to process tens of thousands of MS/MS spectra in a large database, using resources available in most desktop/laptop computers.
Is a self-contained workflow available with a friendly graphical user interface (GUI) with many features available for data analysis and visualization.

However, the previous versions of MiCId’s workflow do not provide protein quantification, AR protein identification, and are only available for the Linux operating system. In this study, we have augmented the MiCId’s workflow to meet the criterion for the identification of AR proteins and we intend to address the other two unmet criteria in the near future. MiCId’s workflow can, however, be used in the Windows operating system via a virtual machine. Details of how to run MiCId’s workflow in the Windows operating system are described in MiCId’s user manual.

The AR protein identification task for an MS/MS workflow can be formulated as follows. First, using data from an MS/MS experiment, a workflow needs to identify the species/strains present in the biological sample. Second, it needs to construct, on the fly, a target protein database to be used for AR protein identifications. Even if a workflow has high sensitivity and high specificity for the identification of microorganisms and proteins, a remaining difficulty to be dealt with in identification of AR proteins is deciding what protein sequences to include in the target protein database. In principle, the ideal target protein database to use would include all the protein sequences obtained directly from the strains present in the biological sample and with AR proteins unambiguously annotated. However, such a database is unobtainable from an MS/MS based proteomics approach, even if strain level identification is attained. It’s standard practice for workflows to use databases such as those hosted by the National Center for Biotechnology Information (NCBI) to obtain protein sequences for as-yet-to-be-identified strains to build a target protein database. A target protein database constructed by using this procedure is an approximation to the ideal target protein database, because the strains present in the biological sample could have gained new proteins via horizontal gene transfer and mutations through rapid multiplication and environmental pressure [50,51]. To mitigate this issue, MiCId constructs on the fly a target protein database made of proteins from the reference/representative proteomes of confidently identified species and AR proteins from a high-quality AR database [27,52,53]. This strategy is employed because the proteomes of reference/representative strains are proteome assemblies of higher quality; hence, they are to be used as anchors for the analysis of closely related proteomes within the same taxonomic group [54]; by including a comprehensive AR protein database in a target protein database, MiCId’s workflow can potentially deal with the horizontal AR gene transfer and potentially can allow the presence of few mutations occurring in the AR proteins to be identified. Overall, the target protein database used in MiCId’s strategy is not too far off from the ideal target database because the proteomes of most strains under a given species share a significant number of highly homologous proteins [55,56] and the inclusion of AR proteins in a general manner takes care of the possible gain, via horizontal gene transfer, of known AR proteins.

We have used five MS/MS datasets, consisting in total of 126 HPLC-MS/MS datafiles (each containing about 20,000 to 30,000 spectra), covering 10 antibiotic resistant bacterial strains, to evaluate the newly augmented MiCId’s workflow in terms of AR protein identifications. In our evaluation, AR proteins are identified at the AR protein family level, following the AR protein family classification used by the CARD database [52,57]. Identification of AR proteins is performed at the family level because of the large number of highly homologous AR proteins within most AR protein families. (Many AR proteins within the same AR family differ from each other by only one to few amino acid residues.) The high degree of protein sequence similarity makes the task of distinguishing among individual proteins beyond the AR protein family level not always possible; especially when data-dependent acquisition mode is used in MS/MS experiments. Although identification of the exact AR protein is not always possible, obtaining identifications at the AR protein family level are enough to improve antibiotic treatments for patients suffering from bacterial infections, since AR proteins within the same AR protein family are largely resistant to the same antibiotics.

In our evaluation, we have shown that MiCId’s workflow has a sensitivity of aproximately 85% (with an estimated lower bound of 72%) and a precision greater than 95% in the identification of AR protein families. We have demonstrated, using an MS/MS dataset from samples of two human pathogens, that MiCId’s workflow can be employed to investigate possible mechanisms of antibiotic resistance in bacteria. We have also shown that MiCId’s workflow can provide microorganismal identification, protein identification, sample biomass estimation, and AR protein identification in 6–17 minutes using computer resources that are available in most desktop and laptop computers. The new MiCId version v.07.01.2021, designed to run in a Linux environment and tested under (i) CentOS Linux release 7.9.2009, (ii) Red Hat Enterprise Linux Server release 7.9, (iii) Ubuntu release 18.04.3, and (iv) Windows 10 using Oracle VirtualBox 6.1.22 running Ubuntu release 18.04.3, is freely available for download at https://www.ncbi.nlm.nih.gov/CBBresearch/Yu/downloads.html.

## 2 Materials and Methods

### 2.1 MiCId’s AR Protein Identification Algorithm

MiCId’s workflow is augmented to allow for AR protein identifications. Mi-CId’s workflow contains procedures for taxonomic identifications, biomass estimations, and protein identifications. In the workflow, it is the protein identification part that gets augmented for the purpose of AR protein identifications. Below, we summarize MiCId’s workflow and highlight the augmentations required. MiCId begins by querying a sample’s MS/MS spectra in the microorganismal database, containing protein sequences from reference and representative genomes, for the identifications of microorganismal peptides; these identified microorganismal peptides are then used for taxonomic identifications, via an iterative approach at each taxonomic level, and for relative taxa biomasses estimates within the sample [29,30]. The proteins from the reference/representative proteomes of species identified with *E*-value ≤ 0.01 and *prior* ≥ 0.01 are then assembled on-the-fly for protein identification.

In the augmented MiCId, we add to the aforementioned protein database AR proteins from an AR database. Namely, in the protein identification procedure, MiCId now queries the updated protein database (combining the protein database constructed on-the-fly and the AR database) with MS/MS spectra to identify peptides for protein and AR protein identifications. (This should not be confused with the peptide identifications needed for taxonomic identifications and biomass estimates.) MiCId uses the scoring function and statistics from the database search tool RAId_DbS [49] to score peptides and for assigning statistical confidences, *E*-values, to identified peptides. Identified peptides are then used as evidences for protein identifications. See Figure 1 for an overview of MiCId’s workflow.

**Fig. 1:**
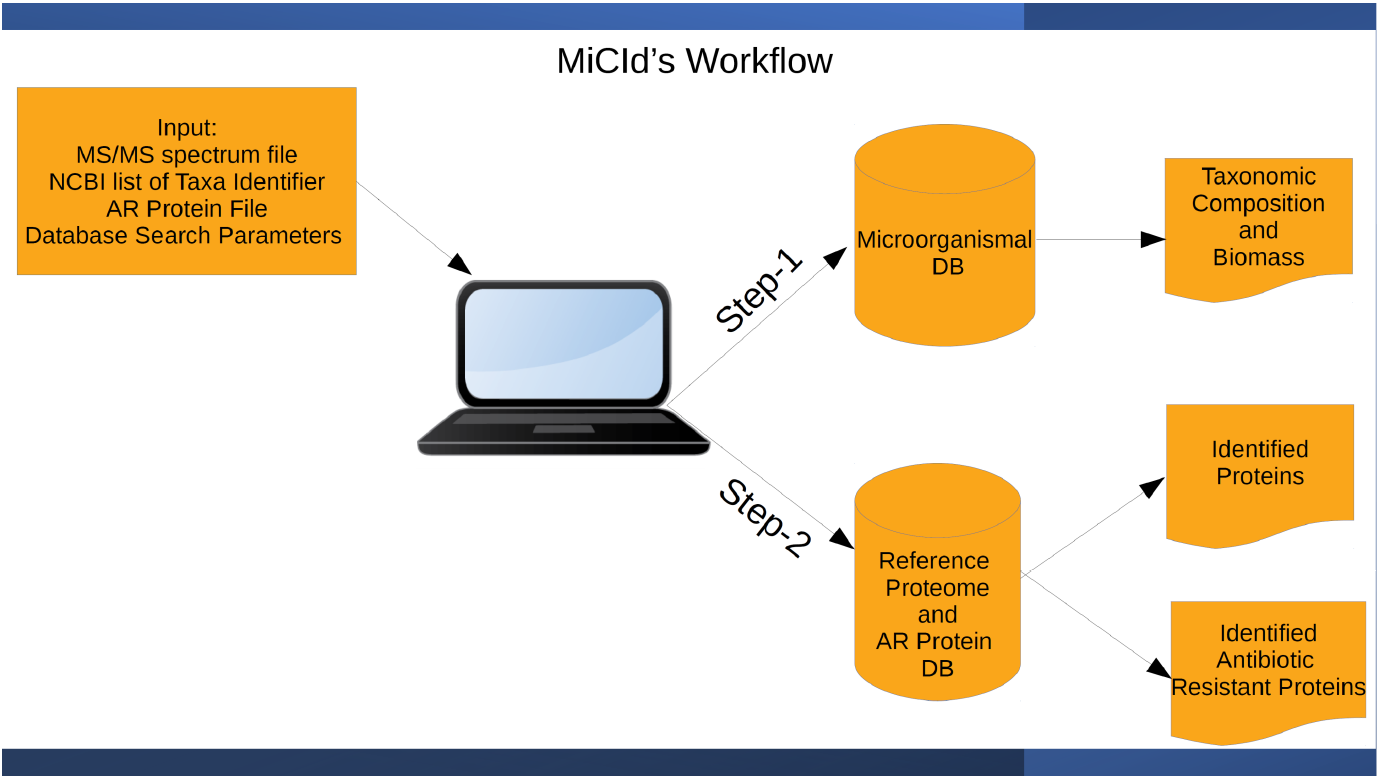
MiCId’s workflow overview. To execute MiCId, users must provide the following input: a list of taxonomic identifiers taken from the NCBI, an experimental datafile (containing MS/MS spectra) of a microorganism sample, an antibiotic resistance (AR) protein file, and the parameters for database search. The list of taxonomic identifiers is used by MiCId to download from the NCBI the Fasta files of the protein sequences for all the taxa specified along with their taxonomic information. The downloaded protein Fasta files and the taxonomic file are used to create the microorganismal database. In step-1, the MS/MS spectra are queried in the microorganismal database in order to determine the taxonomic composition (via an iterative approach that propagate only taxa identified at one level to identifications at the next level) and the relative biomasses of microorganisms in the sample [29, 30]. In step-2, the newly augmented step, MiCId generates a protein database that includes protein sequences from reference/representative strain of species identified with *E*-value ≤ 0.01 and *prior* ≥ 0.01, and from the user specified AR protein file. The MS/MS spectra are then used to query this database to perform protein identifications, AR proteins included.

When performing protein identifications, proteins that share a large number of identified peptides are grouped as a cluster. To control the number of identified proteins, several existing methods [58] report those similar proteins as one. Adopting the same idea, we implemented this approach via two clustering procedure: (1) a peptide-centric clustering procedure and (2) a protein-similarity clustering procedure.

For the peptide-centric clustering procedure, identified proteins are sorted by the number of identified evidence peptides in descending order and the rank of a protein in the sorted list is used as that protein’s cluster index.

The peptide-centric similarity of protein *β* to protein *α* is computed as the ratio of the weighted count of shared identified peptides to the weighted count of identified peptides belonging to protein *β* (with *π_i_* denoting an arbitrary identified peptide)

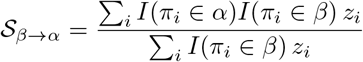

where *I*(*π_i_* ∈ *α*) is an indicator function taking value 1 (0) if *π_i_* is present (absent) in protein *α* and *z_i_* is the weighted count of *π_i_* given by

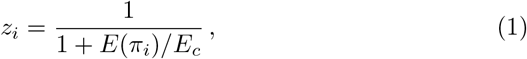

with *E_c_* being the *E*-value cutoff used to control the expected number of false positive peptides at the 5% PFD [59].

One then computes 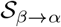 for all protein pairs (*β, α*) with *α* ranking above *β*. Starting with the first protein *γ*_1_ on the sorted list as the reference protein, all other lower-ranking proteins *β* with 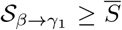 will have their cluster indexes changed to that of the reference protein. One then moves the reference point (from the first) to the second protein denoted by *γ*_2_, all remaining lower-ranking proteins *β* with 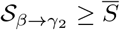 will have their cluster indexes changed to that of the reference protein. The reference point is then moved to the third protein *γ*_3_ and the process continues till the reference point moves through all proteins in the list. The process is iterated for 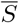 ranging from 1 to 0.5 in steps of 0.025. An exception to the above clustering rule, however, is introduced to properly underscore identified *unique* peptides, each of which is a subsequence to only one protein within the proteins identified. When a protein’s unique evidence peptides’ *E*-values yield a combined *E*-value ≤ *E*_1_ [60], where *E*_1_ is the *E*-value corresponding to having one expected false positive peptide, our method does not allow this protein to be clustered to any other reference proteins. At the end of each iteration, a protein cluster is considered as a protein and the number of unique peptides (to each protein) and their combined *E*-value [60] are recomputed. In the above clustering procedure, a protein that retains its original cluster index is referred to as the head of the cluster, while other proteins in the cluster members of the cluster.

To cluster highly homologous proteins that have not clustered together under the peptide-centric clustering procedure, we next invoke the protein-similarity clustering procedure within which pairwise sequence alignments by BLAST [61] are performed among the heads of protein clusters. The length normalized BLAST bit-score is defined as the BLAST bit score divided by the length of the longer of the two sequences aligned. A protein (or a cluster head) *γ* that has a length normalized BLAST bit-score greater than a threshold with respect to a higher ranking protein and less than 2 unique evidence peptides with *E*-values ≤ *E*_1_ is clustered to that higher ranking protein; hence, *γ* loses its status as a cluster head. This procedure is iterated with the threshold starting from highest length normalized bit-score attainable down to 1.6. The 2 unique evidence peptide rule [58,62] with the number of false positive peptides controlled at 1, *E*-values ≤ *E*_1_, is employed to infer the identification of highly homologous proteins.

Once the proteins have been clustered, a unified *P*-value is computed for all identified proteins via combining evidence peptides’ *E*-values, as previously described [40]. The protein with most significant unified *P*-value within a cluster (containing one or more proteins) is then promoted to be the head of the cluster and the other proteins in the same cluster the members of the cluster (MC). The *P*-value assigned to the head of a cluster is regarded as the *P*-value associated with that cluster.

As a consequence of the clustering procedures employed, two possible scenarios need to be considered when trying to identify AR proteins. In the first scenario, the head of an identified cluster is an AR protein. For this case, no additional step is required, and the identified protein is considered a possible AR protein. The second scenario occurs when the head of an identified protein cluster is not an AR protein. When this happens, in order to decide if the identified protein should be considered a possible AR protein, MiCId employs BLAST [61] and queries the head of the identified protein cluster in the specified AR database. The best BLAST result for each queried protein is kept and only the heads with a length-normalized BLAST bit-score ≥ 1.0 are reported. Protein cluster heads with a length-normalized BLAST bit-score ≥ 1.6 are deemed an AR protein with confidence by MiCId and the question of whether or not to regard the other reported protein cluster heads as possible AR proteins would require further investigations that are beyond the scope of the current manuscript.

MiCId aims to identify AR protein candidates that are globally homologous to the AR proteins already validated (e.g., proteins in an AR database). Therefore, it is important to normalize the BLAST bit-score by the sequence length to properly quantify the quality of the global similarity. Panels A and B of Figure S1 display the histograms for the BLAST bit-scores and for the length-normalized BLAST bit-scores of the self alignment of each of the 3109 proteins taken from the ResFinder database. Comparing the histograms of panels A and B of Figure S1, one finds that the length-normalized BLAST bit-score has a much narrower score range, [1.6, 2.1], than the BLAST bitscore, [100, 1800], indicating a better choice of scores for learning a score cutoff value. While the BLAST bit-score has been shown to be more discriminative than the BLAST *E*-values in capturing functional homologous sequences [52], it does not scale with the length of the sequences aligned. The BLAST bitscore provides information about the quality of the local similarity between two sequences, not the global similarity [63]. The length-normalized BLAST bit-score cutoff value of 1.6 for designating an identified protein an AR protein was learned from the PFD curves generated with different cutoff values. Figure S2 shows the various PFD curves. The PFD curves generated with cutoff values ≥ 1.6 have comparable performance. To ensure that no AR protein is missed, the cutoff value of 1.6 is chosen to cover the entire score range obtained for the self alignment of all proteins in the ResFinder database.

### 2.2 MS/MS datasets

A total of five MS/MS datasets were used for this study. One dataset, generated in-house PXD026634, is composed of 21 experimental MS/MS datafiles from samples of five bacteria strains. The other four datasets, downloaded from the ProteomeXchange Database (PD) [64], contain 105 experimental MS/MS datafiles from samples of five other bacterial strains. For seven bacterial strains used in this study, one may download their complete genomic sequences [24, 65–67] and protein sequences from the National Center for Biotechnology Information (NCBI) databases [68]. In Table S1, we provide the pertinent information for each MS/MS dataset.

#### 2.2.1 In-house MS/MS dataset

##### 2.2.1.1 Bacterial Strains

Two carbapenem-resistant *P. aeruginosa* strains were included in the study. Strain CCUG 51971 (= PA 66) was isolated from a human urine sample, at the Karolinska Hospital (Stockholm, Sweden), carrying OXA-35, OXA-488, PDC-35 and VIM-4 [65]. The VIM-4 metallo-*β*-lactamase is responsible for the high carbapenem resistance levels (Minimum inhibitory concentration (MIC) of imipenem and meropenem greater than 256 *μ*g/ml; MIC of imipenem + ethylenediaminetetraacetic acid [EDTA] = 6 *μ*g/ml) [65]. Strain CCUG 70744 was isolated from a human sputum sample, at the Sahlgrenska University Hospital (Gothenburg, Sweden), carrying OXA-905 and PDC-8 [67,69,70].

Furthermore, one *E. coli* and two *K. pneumoniae* strains, isolated from various clinical samples at the Sahlgrenska University Hospital, carrying different *β*-lactamases (including extended spectrum *β*-lactamases, ESBL, and carbapenem resistance genes) were included in the study. *E. coli* CCUG 70745 isolated from human feces, carrying CMY-6, CTX-M-15, NDM-7,and OXA-1; *K. pneumoniae* CCUG 70742, isolated from human urine carrying CTX-M-15, OXA-1, OXA-48 and TEM-1; and *K. pneumoniae* CCUG 70747, isolated from human wound, carrying KPC-2, SHV-200, TEM-1 and VIM-1 [67]. Lyophiles of all strains were obtained from the Culture Collection of University of Gothenburg (CCUG, Gothenburg, Sweden; www.ccug.se). The strains were reconstituted on Müller-Hilton agar (Substrate Unit, Department of Clinical Microbiology, Sahlgrenska University Hospital), at 37°C, for 24 h.

##### 2.2.1.2 Cultivation conditions

Preinocula were made in 4 ml of Müller-Hilton broth (MHB), incubated, at 37°C, overnight, with orbital shaking (250 rpm). Subsequently, the *P. aeruginosa* strains were cultivated in 50 ml of MHB, with a final inoculum size of 5 x 10^5^ colony forming units (CFU) ml-1, in baffled Erlenmeyer flasks of 250 ml capacity, at 37°C, for 20 h, with orbital shaking (250 rpm). Strain CCUG 70744 was cultivated with 0, 2, 4 and 8 *μ*g/ml of meropenem. Strain CCUG 51971 was cultivated with 0, 8, 128 and 256 *μ*g/ml of meropenem. For *E. coli* CCUG 70745, *K. pneumoniae* CCUG 70742 and *K. pneumoniae* CCUG 70747 strains, the bacterial biomass was adjusted to McFarland 0.5 by diluting the culture using phosphate-buffered saline (PBS). 100 *μ*l of the McFarland 0.5 was diluted with 1900 *μ*l of PBS (dil. 1:20). 400 *μ*l of the bacterial suspension was added to glass tubes and MHB was added together with the antibiotic (AB; ertapenem) from a 10 mg/ml stock solution of (or in the case of no antibiotics condition, additional broth) to a final volume of 4.4 ml. For *E. coli* CCUG 70745 = 3975 *μ*l of MHB + 25 *μ*l AB + 400 *μ*l bacterial suspension (final AB concentration 56 *μ*g/ml), for *K. pneumoniae* CCUG 70742 = 3990.5 *μ*l MHB + 9.5 *μ*l AB + 400 *μ*l bacterial suspension (final AB concentration 21 *μ*g/ml) and for *K. pneumoniae* CCUG 70747 = 3987.5 *μ*l MHB + 12.5 *μ*l AB + 400 *μ*l bacterial suspension (final AB concentration 28 *μ*g/ml). The strains were cultured for 16 to 20 hours with shaking (250 rpm) at 37°C.

##### 2.2.1.3 Sample preparation

Bacterial biomass was collected and suspended in PBS. Bacterial cell suspension optical densities (OD) were measured at a wavelength of 600 nm and adjusted in 1 ml PBS to OD_600_ = 0.8 (10^9^ CFU/ml). The bacterial biomass was washed with PBS three times by centrifuging the sample for 5 min at 12,000 × g, discarding the supernatant and resuspending the pellet in 1.0 ml PBS. Finally, cells were resuspended in 150 *μ*l of PBS and transferred to small vials (200 *μ*l) containing glass beads (G1145, Sigma-Aldrich, St Louis, MO, USA), in preparation for cell lysis, by bead-beating, using a TissueLyser (Qiagen, Hilden, Germany; settings: frequency 25 Hz for 5 min). The bacterial lysates were frozen at −20 °C until analysis. For digestion of proteins, to generate peptides, sodium deoxycholate (SDC, 5% in 20 mM ammonium bicarbonate, pH 8) was added to 1% (w/v) final concentration. Trypsin (2 *μ*g/ml, 100 *μ*l ammonium bicarbonate, 20 mM pH 8) was added and samples were digested for approximately 8 h at 37°C. SDC was removed by precipitation by addition of formic acid (FA) followed by centrifugation at 13,000 × g for 10 min. Supernatants containing the peptides were stored at −20 °C until analysis. The peptide samples were not reduced or alkylated *prior* to MS analysis.

##### 2.2.1.4 Liquid chromatography - tandem mass spectrometry (LC–MS/MS) acquisition

Samples were desalted. The samples and fractions were dried and reconstituted in 3% acetonitrile, 0.2% formic acid. Samples were analyzed on a QExactive HF mass spectrometer interfaced with Easy-*η*LC 1200 liquid chromatography system (Thermo Fisher Scientific). Peptides were trapped on an Acclaim Pepmap 100 C18 trap column (100 *μ*m x 2 cm, particle size 5 *μ*m, Thermo Fischer Scientific) and separated on an in-house packed analytical column (75 *μ*m x 30 cm, particle size 3 *μ*m, Reprosil-Pur C18, Dr. Maisch) using a gradient from 5% to 80% acetonitrile in 0.2% formic acid over 90 min at a flow of 300 *η*L/min. The instrument operated in data-dependent mode where the precursor ion mass spectra were acquired at a resolution of 60 000, m/z range 400-1600. The 10 most intense ions with charge states 2 to 4 were selected for fragmentation using HCD at collision energy settings of 28. The isolation window was set to 1.2 Da and dynamic exclusion to 20 s and 10 ppm. MS2 spectra were recorded at a resolution of 30 000 with maximum injection time set to 110 ms. The mass spectrometry proteomics data have been deposited to the ProteomeXchange Consortium via the PRIDE [71] partner repository with the dataset identifier PXD026634.

#### 2.2.2 Downloaded MS/MS datasets

Four publicly available datasets, previously used in two different studies on the identification of AR proteins in bacteria, were downloaded from the ProteomeXchange Database (PXD) (http://www.proteomexchange.org/). Dataset PXD004321 was taken from the study of the computational method TCUP on the identification of AR proteins [24]. This dataset contains 6 experimental MS/MS datafiles from samples of a ESBL *E. coli* strain CCUG 62462, carrying CTX-M-15 and TEM-1 [66], the CCUG 62462 strain was grown in pure cultures without and with cefotaxime at 1000 *μ*g/ml. Dataset PXD011105, containing 35 experimental MS/MS datafiles, was taken from the study on the mechanism of antibiotic resistance of two clonal isolates (the *P. aeruginosa* strain CLJ1 antibiotic-sensitive isolate and the *P. aeruginosa* strain CLJ3 multidrug-resistant isolate obtained from the same patient at different times) grown in pure cultures with carbenicillin at 200 *μ*g/ml [72]. Dataset PXD005587, containing 24 experimental MS/MS datafiles, was taken from the investigation on proteomics changes due to antibiotic-dependent perturbations in ESBL *K. pneumoniae* strain 34618, grown in pure cultures without and with doxycycline or streptomycin [31]. Dataset PXD010244, containing 40 MS/MS datafiles, was taken from the research on the mechanism of antibiotic resistance in ESBL *K. pneumoniae strain* KpV513, grown in pure cultures without and with doxycycline or streptomycin or doxycycline and streptomycin [32].

### 2.3 MS/MS data analysis using MiCId workflow

All datasets were analyzed using the MiCId (v.07.01.2021) workflow [21,29, 30]. The peptide-centric microorganismal database used for analysis includes all reference and representative proteomes of bacteria (12,703 strains in total) that are available in the National Center for Biotechnology Information (NCBI) database as of February 4, 2021. The reference proteome of one *Homosapiens* is also included for two reasons. First, human proteins are a major component of microorganism samples when obtained directly from human host. Second, human proteins (mostly keratin) are also frequently identified, albeit at lower abundances, even in microorganism samples from laboratory cultures. The reference proteomes of *Bos taurus* and *Saccharomyces cerevisiae* are also included because they are present respectively in the Mueller Hinton Broth and in the Luria-Bertani Broth; both broth media are routinely used to grow bacterial cultures. The protein sequence Fasta files for the 12,703 organisms along with the file containing taxonomic information were downloaded from the NCBI database at ftp://ftp.ncbi.nlm.nih.gov/genomes/Bacteria and at https://www.ncbi.nlm.nih.gov/Taxonomy on February 4, 2021. In total, 60,176,722 protein sequences were downloaded. As previously described [29], to speed-up MS/MS spectrum analysis, MiCId processes the protein sequences and the taxonomic file into a peptide-centric microorganismal database. The final size of the peptide-centric microorganismal database is 100 GB.

To allow for AR protein identifications, in addition to the database mentioned above, MiCId included in its search scope AR proteins from one of the following databases: ResFinder [27], CARD [52] or NDARO [53, 73]. Users also have the option to provide for MiCId their own assembled AR protein database in a Fasta file. Table S2 lists the protein identifiers for the AR proteins along with the taxonomy identifiers and scientific names of the organisms included in MiCId’s databases.

While querying the database with PXD004321, PXD026634, PXD011105, PXD005587, and PXD010244 datasets, the following search parameters were employed. The digestion rules of trypsin and lys-c were assumed with up to two missed cleavage sites per peptide. The mass error tolerance of 5 ppm was set for the precursor ions and 20 ppm for the product ions except when analyzing PXD011105, PXD005587, and PXD010244 (10 ppm for the precursor ions). For PXD004321 and PXD026634, cysteines were unmodified and for PXD011105, PXD005587, and PXD010244 iodoacetamide was used as the reduction agent, changing the molecular mass of every cysteine from 103.00919 to 160.030647 Da. RAId’s Rscore, using b and y ions as evidence, was used to score peptides. The statistical significance of each peptide was assigned via RAId_DbS’s theoretically derived peptide score distribution [49]. The largest (cutoff) *E*-value for a peptide to be reported was set to 1. For taxa identifications at the genus level and lower, all microorganisms under the genus *Shigella* were excluded from consideration to avoid classification ambiguity because some researchers have argued that taxonomically *Shigella* should be classified under *Escherichia coli* [74].

### 2.4 Protein identification using Proteome Discoverer software

The MS/MS datasets PXD026634 and PXD004321 (containing a total of 27 MS/MS experimental datafiles from 6 antibiotic resistant bacterial strains), used to estimate the sensitivity of AR protein identifications via MiCId’s Workflow, were also analyzed via Proteome Discoverer version 2.4 (Thermo Fisher Scientific). Protein identification using Proteome Discoverer version 2.4 was performed using Mascot search engine v. 2.5.1 (Matrix Science, London, UK). The database searched was generated from the genomes of the strains included in the study, i.e., containing only genomes from the TP strains. The precursor ion (MS^1^) mass error tolerance is set to 5 ppm while the fragment ions’ mass error tolerance is set to 30 mDa. Tryptic peptides were accepted with up to 1 missed cleavage and methionine oxidation was set as a variable modification. Target decoy was used for peptide-to-spectrum match (PSM) validation with the strict false discovery rate (FDR) threshold of 1% and minimum Mascot Score of 15.

## 3 Results and Discussion

In this section we present the evaluation of using MiCId’s workflow to identify AR proteins. First, we use 126 experimental MS/MS datafiles to assess MiCId’s AR protein identification strategy. Second, we estimate the sensitivity of AR protein identifications via MiCId using 27 experimental MS/MS datafiles from samples of 6 antibiotic resistant bacteria strains (from 3 species included in the pathogen priority list of the World Health Organization [75] for antibiotics research and development), cultured with and without antibiotics. Third, using 35 experimental MS/MS datafiles from samples of two human pathogens, we employ MiCId’s workflow to investigate possible mechanisms of antibiotic resistance.

Following the AR protein classification used by the CARD database [52, 57], one finds that there are large numbers of highly homologous AR proteins within most AR protein families and expects this family level crowdedness to remain as AR protein databases continue to grow. As an illustration, we note that each AR protein within the *β*-lactamase family has very homologous sequences within the family: if one takes an AR protein as the query to align with each of the rest of the AR proteins in the *β*-lactamase family, for the best pairwise alignment, there are many high score alignments and the best of which has an average length normalized BLAST bit-score ≥ 2. This is shown in Table S3. The high degree of similarity for AR proteins in the same family makes distinguishing among AR proteins at finer-than-family level not always possible particularly when data acquisition in the MS/MS experiment is untargeted. For these reasons, during our evaluation, identified AR proteins are counted as true positives and false positives at the AR protein family level. For example, assume a bacteria strain contains OXA-1 and OXA-48 proteins, leading to 2 proteins in the OXA family; if during the analysis MiCId identifies 3 OXA proteins, then the 2 best ranking OXA proteins identified are counted as true positives and the remaining OXA protein is counted as a false positive. For some AR protein families that are not yet overly represented/crowded in the database, correct identification can be achieved at finer-than-family when closely related homologous proteins are present in the database. For example, for AR proteins from aminoglycoside families, families which are not overly represented in the database, we observed correct identifications for these AR proteins not only at the family level but also at the isoenzyme level [76,77], which is a finer level than the family level. Of course, if the target family (to which the query protein belongs) is too much under-represented in the database, either no identification is made, or misidentifications of the AR protein families occur.

### 3.1 Evaluation of MiCId’s AR Protein Identification Strategy

MiCId’s strategy for AR protein identifications is to first identify species in a microorganismal database and then identify AR proteins in a target protein database composed of proteins from the reference/representative proteomes of confidently identified species and AR proteins from a high-quality AR database [27,52,53]. MiCId’s strategy capitalizes on microorganismal identifications at the species level because high confidence microorganism identifications at taxonomic levels lower than species become challenging due to the lack of discriminative peptides among the ones identified [30,78] when using routinely employed high resolution data-dependent acquisition mode in the MS/MS experiment [79]. In principle, more advanced MS/MS experiments such as targeted MS/MS using selected reaction monitoring (SRM) or parallel reaction monitoring (PRM) can be used for taxonomic identification below the species level by targeting peptides that are unique to taxa at lower taxonomic levels [78,80–83]. However, a limitation of such approaches is that they can only be employed for the identifications of a microorganism within a small, predetermined set of microorganisms. Another reason for employing MiCId’s strategy has to do with the trustworthiness in annotation of the taxonomic database for taxonomic levels below species [84–87]. It is important to mention that although for this study we only included the proteins from strains that are labeled as reference and representative in the microorganismal database, as these are proteins from higher-quality genomes [54,88,89], MiCId’s workflow is not limited to microorganismal database composed of only reference and representative genomes and it can perform microorganismal identifications beyond the species level.

When, for the purpose of protein identifications, selecting a proteome as the representative for a confidently identified species, MiCId relies on a heuristic because under a given species there could be many strains and priority for each has to be established. The heuristic gives strains that are labeled as *reference* first priority and *representative* second priority. Information about reference and representative strains is taken from the RefSeq and GenBank assembly summary files downloaded from the NCBI. If there is more than one reference strain or representative strain for a given species, the strain with the larger number of proteins is selected. When a species has neither reference strain nor representative strain, the proteome from the strain, under that species, with the larger number of proteins is selected. The rule of assigning high priority to the proteomes from reference strains and representative strains is applied because these are proteome assemblies of higher quality and importance that have been curated by the NCBI staff and are to be used as anchors for the analysis of closely related proteomes within the same taxonomic group [54].

For identifications of AR proteins, the ideal target protein database would include all the protein sequences obtained directly from the strains present in the biological sample and with AR proteins unambiguously annotated. From an MS/MS based proteomics viewpoint, such a database is unattainable even if strain level identification is achieved. MS/MS based proteomics approaches relies on databases such as the ones at the NCBI to obtain protein sequences for yet-to-be-identified strains. A target protein database constructed using this procedure would still be an approximation to the ideal target protein database because the strains present in the biological sample could have acquired new proteins via horizontal gene transfer and mutations through rapid multiplication and environmental pressure [50, 51]. By including a comprehensive AR protein database in the target database, MiCId can potentially deal with the horizontal AR gene transfer; through the clustering procedure, MiCId can potentially allow the presence of few mutations occurred in the AR proteins to be identified. Overall, the target protein database used in MiCId’s strategy is not a bad approximation to the ideal target database because the proteomes for most strains under a given species shared a significant number of homologous proteins [55,56] and it includes AR proteins in a global manner, covering the possible acquisition, via horizontal gene transfer, of known AR proteins.

To evaluate MiCId’s strategy for identifying AR proteins, we prepared two sample-specific target protein databases and query them with the same MS/MS datafiles from specific samples. The first target protein database is composed of proteins from the reference proteome of the species present in the sample and AR proteins from the ResFinder database, referred to here as reference strains plus AR database (RA). The other target protein database is composed of proteins from the proteome of the true strain present in the sample and AR proteins from the ResFinder database, referred to here as correct strain plus AR database (CA). Table S4 contains the protein identifiers for each protein used to generate both versions of the sample-specific target protein databases. Plotted in panels A and B of Figure 2 are the PFD curves from querying the RA and CA databases with 62 experimental MS/MS datafiles from samples of seven strains. There are in total 14 target protein databases, 2 for each of the 7 strains that have complete genome sequence available, used in generating panels A and B of Figure 2. The PFD curves in panel A of Figure 2 shows that using the target protein databases RA and CA produced comparable PFD curves. Furthermore, the curves in panel B of of Figure 2 show that using RA and CA databases yield 131 AR protein identifications in common. What wasn’t shown is that there are 12 AR protein identifications, covering 5 AR protein families, not shared: there are 8 PDC protein family identifications and the 1 ant(3”) family identification present in the list identified using the CA database but absent from that using the RA database; on the other hand, 1 TEM family identification, 1 OXA family identification, and 1 ARR family identifications are only found using the RA database. Multiple PDC family identifications are found using both databases: 23 identifications using the CA database and 15 identifications using the RA database. The identification rate of PDC protein family in the CA database is higher because it contains the correct PDC protein PTC38756.1, that belongs to CLJ1 strain, even though this protein is not yet included in the ResFinder database. Also, in the ResFinder database the PDC family –containing only 4 PDC proteins: AAM08942.1, ACQ82815.1, ACQ82807.1, and AAM08945.1– is under-represented, making it difficult to identify the correct PDC using the ResFinder database since even the most homologous PDC protein (AAM08942.1) and the correct PDC protein (PTC38756.1) differ by more than 50 amino acid residues. The discrepancy in true positives in the other 4 AR protein families is also mainly caused by composition difference of the 2 target databases. Table S5 contains pertinent information of all the identified proteins/families in both databases. Panel C of Figure 2 shows that there are 180 AR protein family identifications at the 5% PFD level, when analyzing all the 126 MS/MS data files. Panel D of Figure 2 shows that using a *E*-value cutoff of 0.01 the identification of AR proteins can be controlled at the 5% PFD level. Based on this result, in order to control the false positives at around the 5% PFD level, only AR protein family identifications with *E*-value below 0.01 are deemed true positive with high confidence by MiCId. Table S6 has the list of all identified proteins for all the 126 MS/MS data files.

**Fig. 2:**
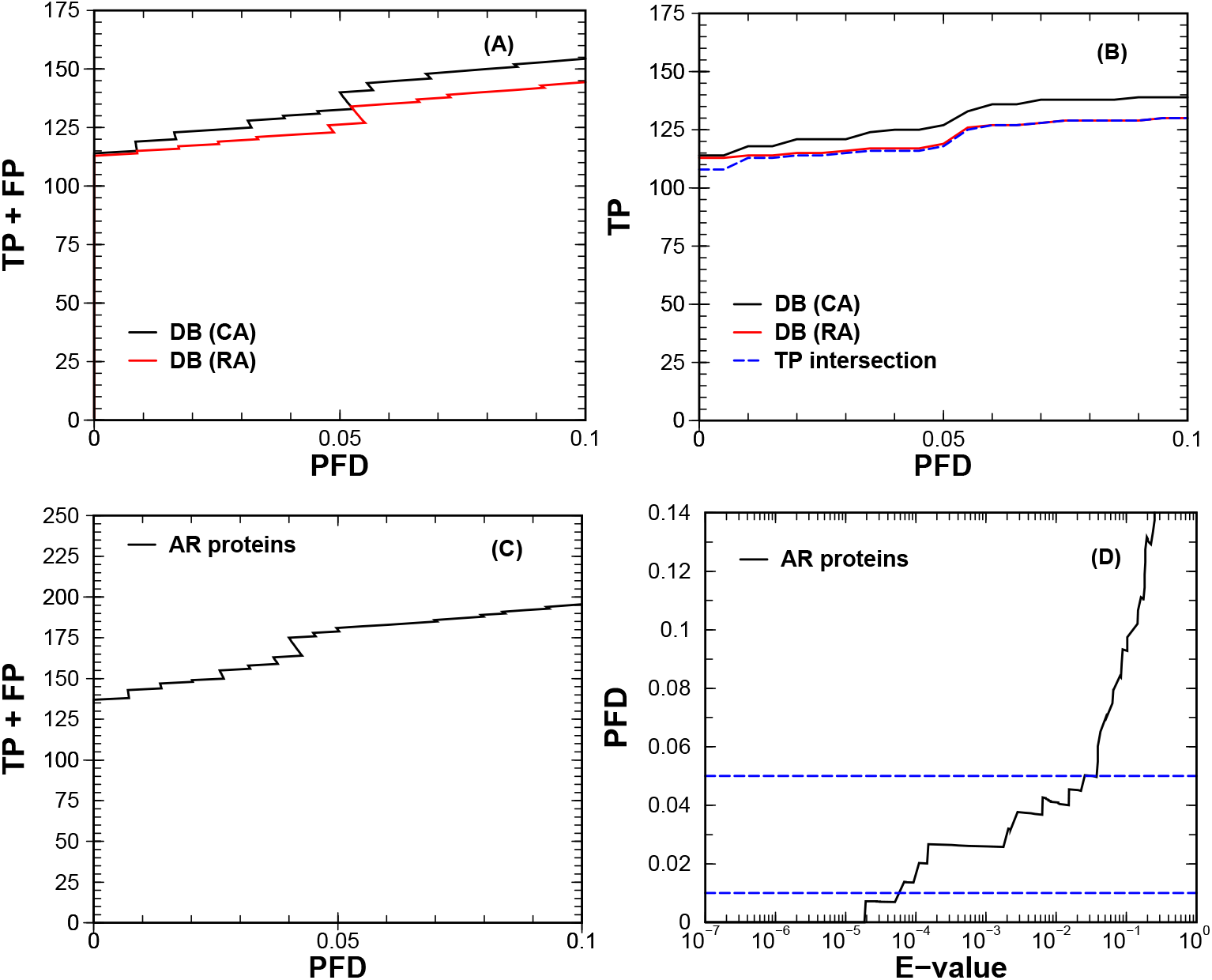
MiCId’s workflow evaluation. Let us mention here again that the abbreviations CA and RA refer respectively to target databases composed of proteins from correct strain plus the chosen AR database and from reference strains plus the chosen AR database. Panels A and B display PFD curves when querying the CA and RA with 62 experimental MS/MS datafiles. Panel A shows that the PFD curves from searching in CA and RA are comparable. Panel B shows that there are 131 true positive antibiotic resistance (AR) proteins identified in common in CA and RA. The PFD values in panels C and D were obtained from querying RA with 126 experimental MS/MS datafiles. Panel C also indicates that 180 AR proteins are identified at the 5% PFD level. Panel D shows that using an *E*-value cutoff of 0.01 the identification of AR proteins can be controlled at the PFD level smaller than 5%. The abbreviations TP, FP and PFD refer respectively to true positive, false positive and proportion of false discoveries.

As mentioned above, a requirement for MiCId’s strategy to work is that it must have accurate species level identification. MiCId achieves accurate microorganism identification with trustworthy confidence assignments by properly computing for every identified taxon an *E*-value and a *prior* probability [29, 30]. For a quality score *S*, the *E*-value reflects the expected number of random taxa with scores the same as or better than *S* [40]. A taxon’s *prior* probability is the probability for an identified taxon to emit any evidence peptide which can also be viewed as that taxon’s protein biomass up to an overall proportionality constant as described earlier [30]. Therefore, identified taxa with small *E*-values and large priors are more likely to be present in the sample. As we have demonstrated, MiCId can control the PFD below 5% by calling true positives only identified taxa with *E*-values ≤ 0.01 and *prior* ≥ 0.01 [29, 30]. Also, MiCId employs an iterative approach for taxa identification at each taxonomic level; only taxa identified at the upper taxonomic level are considered for the next level identifications [29, 30]. As shown in Figure 3, when considering all identifications with *E*-values ≤ 1, MiCId identifies for each of the 126 samples the correct species with only 13 false positives overall. Interestingly, MiCId also identifies *H. sapiens* in 103 samples, *B. taurus* in 18 samples, and *S. cerevisiae* in 12 samples. *H. sapiens* are identified using evidence peptides from keratin proteins detected in the samples. Keratin proteins are a common contaminant to mass spectrometry experiments, usually originating from skin, hair as well as dust, clothing and latex gloves. The identification of *B. taurus* and *S. cerevisiae* in some of the samples is expected as they are present in the broth medium used to grow the bacterial cultures.

**Fig. 3:**
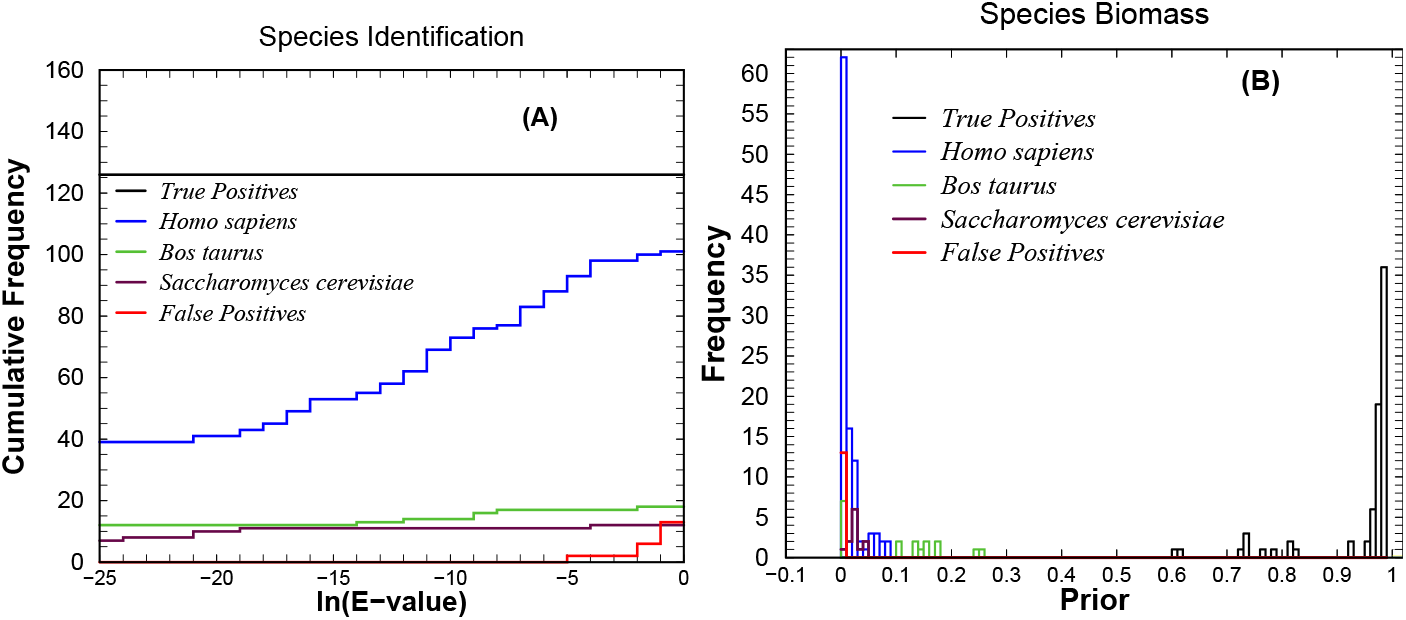
Species level composition for samples 1-126. Plotted in panel A are the cumulative frequency of the number of true positives species, *H. sapiens, B. taurus, S. cerevisiae* and false positives species versus the natural logarithm of the *E*-value of all species identified with *E*-value ≤ 1. Plotted in panel B are the histograms of priors for the true positive species, *H. sapiens, B. taurus, S. cerevisiae* and false positive species. Among the true positives: *E. coli* is identified 10 times, *K. pneumoniae* 72 times and *P. aeruginosa* 44 times. Among the 13 false positives: *Algisphaera agarilytica* is identified 1 time, *Cerasicoccus arenae* 4 times, *Chlorobium tepidum* 2 times, *Desulfosporosinus orientis* 1 time, *Fervidobacterium thailandense* 1 time, *Ktedonosporobacter rubrisoli* 1 time, *Streptococcus thermophilus* 1 time and *Thiofilum flexile* 2 times. *H. sapiens* is identified 103 times, *B. taurus* 18 times, and *S. cerevisiae* 12 times. To control the proportion of false discoveries below 5% only species identified with *E*-value ≤ 0.01 (ln(*E*-value) = −4.6) and *prior* ≥ 0.01 are considered true positives with high confidence. When employing the recommended cutoffs of *E*-value ≤ 0.01 and *prior* ≥ 0.01 MiCId still identifies all true positives with no false positives.

When imposing the recommended cutoffs, *E*-values ≤ 0.01 and *prior* ≥ 0.01, to control the PFD below 5%, MiCId still identifies correctly the true positive species out of each of the 126 samples. This is because, as shown in Figure 3, all the true positives are identified with much lower *E*-value than 0.01 and much larger *prior* than 0.01. However, with the recommended cutoffs, *H. sapiens* is identified now in 41 samples, *B. taurus* in 11 samples, *S. cerevisiae* in 11 samples, and no false positives. In terms of the *prior*, reflecting the taxon’s relative protein biomass, one would expect it to be very close to 1 for true positive species identified, given that the samples each is assumed to contain a single microorganism. The main reason that it deviates from 1 is because out of the 126 samples, when one only imposes *E*-values ≤ 1 for reporting identification, 105 samples has, in addition to the underlying microbe, identifications matching some of the following three organisms: *H. sapiens*, *B. taurus* and *S. cerevisiae*. For these 105 samples, as shown in panel B of Figure 3, *H. sapiens* contributes to the overall protein biomasses with *prior* values ranging from 0.00076 to 0.085 with the average value 0.016; *B. taurus* has *prior* values ranging from 0.0019 to 0.25 with the average value 0.1; *S. cerevisiae* has *prior* values ranging from 0.0011 to 0.048 with the average value 0.026. These nonzero *prior* values for *H. sapiens*, *B. taurus*, and *S. cerevisiae* cause the observed deviation of the *prior* value from 1 for the TP. Table S7 has the pertinent information of the identified species for each sample.

It is important to mention that the taxa identification results reported by MiCId are not filtered by using the recommended cutoff to avoid incidental false negatives. MiCId reports the complete list of identified taxa using a color-coded scheme. Identified taxa passing the recommended cutoffs, *E*-values ≤ 0.01 and *prior* ≥ 0.01, are highlighted in green for high-confidence in being a true positive; taxa identified with *E*-value ≤ 1 and *prior* ≥ 0.001, are highlighted in yellow for low-confidence in being a true positive; and taxa identified with *E*-value >1 or *prior* < 0.001 are highlighted in red for no-confidence in being a true positive.

### 3.2 Estimate for Sensitivity of AR protein identifications via MiCId’s Workflow

Having computational methods that can correctly identify bacteria and also their AR proteins is among the most important research fronts for fighting infections. We demonstrate the usefulness of MiCId’s workflow in serving as such a computational method in this subsection and next. We use datafiles from some bacteria containing *β*-lactamase proteins as examples for reasons listed below. First, *β*-lactam antibiotics are the most prescribed class of antibiotic to fight infections globally [90]; second, in the United States, about 65% of the antibiotics prescribed are *β*-lactam antibiotics [91]. Of special importance in this class of antibiotic are carbapenems. Carbapenems have a broad spectrum of activity and are usually used as the last-line of the defense for seriously ill patients suspected of harboring resistant bacteria [92]. Evidently, correct identifications of carbapenem resistance can help significantly in fighting infections. Also, *β*-lactamase proteins can be harbored by plasmid and when this occurs they can be easily transmitted into different bacteria cells, introducing resistance to the bacteria [90,93,94].

To estimate the sensitivity of MiCId’s workflow on the identification of AR proteins, we used 27 MS/MS experimental datafiles from 6 antibiotic resistant bacterial strains (from 3 species included in the pathogen priority list of the World Health Organization [75] for antibiotics research and development), cultured with and without *β*-lactam antibiotics. The three *β*-lactam antibiotics used belong to two classes of antibiotics: belonging to the cephalosporin class is cefotaxime, and belonging to the carbapenem class are ertapenem and meropenem. Each of the bacterial strains carries between two to four predicted *β*-lactamase proteins and shows resistance to a variety of antibiotics [65–67,70, 95]. *β*-lactamase proteins for these strains were computationally predicted using ResFinder [27]. Table S8 provides a protein-centric view. This table lists for each predicted *β*-lactamase the strains contain it and the names of *β*-lactam drug classes it resists. For the purpose of estimating the sensitivity value, we view each possible *β*-lactamase identification per experiment as a different event. Summing the numbers of possibly identifiable *β*-lactamase proteins from each of the 27 experiments, one obtains a total of 88 potential true positives. This may be viewed as the maximum set of the true positives. An avid reader may ask what happens if some AR proteins, in this case *β*-lactamase proteins, are missed from the database. When that happens, because these proteins will never be identified, they do not contribute counts to either the numerator or the denominator while the sensitivity value is computed. Hence, for the purpose of assessing the sensitivity, one does not need to worry about AR proteins that are not included in the database. On the other hand, a predicted AR protein may never be observed because it is usually expressed in low abundance or it is not even a true protein in the corresponding microorganism’s proteome. When this is the case, it becomes inappropriate to use the maximum TP set as the TP set for the purpose of estimating the sensitivity value.

When using all the 88 possible identifications as the TP set, one obtains a sensitivity value of 72.7% (64/88). This may be viewed as the lower bound of the sensitivity of MiCId’s workflow. If one excludes from the TP set the *β*-lactamases—OXA-1 in *K. pneumoniae* CCUG 70742, OXA-488 in *P. aeruginosa* CCUG 51971, and OXA-905 in *P. aeruginosa* CCUG 70744—that were never confidently observed in any of the corresponding experiments considered, one obtains a sensitivity value of 85.3% (64/75). This sensitivity value may be viewed as the typical sensitivity value while employing MiCId’s workflow.

Table 1 shows the identification results of *β*-lactamase protein families for all the 27 MS/MS experiments. Displayed in Table 1 are: 64 identifications with *E*-value ≤ 0.01 highlighted in green and marked with a ✔, 6 identifications with 0.01 < *E*-value ≤ 1 highlighted in yellow and marked with a ✔, 5 cases of missed identification (while identified in other samples) marked with a ✘, and 12 cases of no identification with no marks.

**Table 1:** Identification results of *β*-lactamase proteins from culture samples of six antibiotic resistant strains cultivated with and without *β*-lactam antibiotics

| *E. coli* CCUG 62462 |  |  |  |  |  |  |
| --- | --- | --- | --- | --- | --- | --- |
| AR Protein | SN-1<br>NA | SN-2<br>NA | SN-3<br>NA | SN-4<br>CTX<br>1mg/ml | SN-5<br>CTX<br>1mg/ml | SN-6<br>CTX<br>1mg/ml |
| CTX-15 | ✓ | ✓ | ✓ | ✓ | ✓ | ✓ |
| TEM-1 | ✓ | ✓ | ✓ | ✓ | ✓ | ✓ |
| *E. coli* CCUG 70745 |  |  |  |  |  |  |
| AR Protein | SN-7<br>ETP<br>56μg/ml | SN-8<br>ETP<br>56μg/ml | SN-9<br>NA | SN-10<br>NA | | |
| CTX-15 | ✓ | ✓ | ✓ | ✓ |  |  |
| CMY-6 | ✓ | ✓ | ✓ | ✓ |  |  |
| NDM-7 | ✓ | ✓ | ✓ | ✓ |  |  |
| OXA -1 | ✓ | ✓ | ✓ | ✓ |  |  |
| *P. aeruginosa* CCUG 51971 |  |  |  |  |  |  |
| AR Protein | SN-11<br>NA | SN-12<br>MEM<br>8μg/ml | SN-13<br>MEM<br>32μg/ml | SN-14<br>MEM<br>128μg/ml | SN-15<br>MEM<br>256μg/ml | |
| OXA-35 | ✓ | ✓ | ✓ | ✓ | ✓ |  |
| OXA-488 | ✓ | ✗ | ✗ | ✗ | ✗ |  |
| VIM-4 | ✓ | ✓ | ✓ | ✓ | ✓ |  |
| PDC-35 | ✗ | ✗ | ✗ | ✓ | ✓ |  |
| *P. aeruginosa* CCUG 70744 |  |  |  |  |  |  |
| AR Protein | SN-16<br>NA | SN-17<br>MEM<br>2μg/ml | SN-18<br>MEM<br>4μg/ml | SN-19<br>MEM<br>8μg/ml | | |
| PDC-8 | ✓ | ✓ | ✓ | ✓ |  |  |
| OXA-905 |  |  |  |  |  |  |
| *K. pneumoniae* CCUG 70742 |  |  |  |  |  |  |
| AR Protein | SN-20<br>ETP<br>21μg/ml | SN-21<br>ETP<br>21μg/ml | SN-22<br>ETP<br>NA | SN-23<br>ETP<br>NA | | |
| CTX-15 | ✓ | ✓ | ✓ | ✓ |  |  |
| OXA-1 |  |  |  |  |  |  |
| OXA-48 | ✓ | ✓ | ✓ | ✗ |  |  |
| TEM-1 | ✓ | ✓ | ✓ | ✓ |  |  |
| *K. pneumoniae* CCUG 70747 |  |  |  |  |  |  |
| AR Protein | SN-24<br>ETP<br>28μg/ml | SN-25<br>ETP<br>28μg/ml | SN-26<br>ETP<br>NA | SN-27<br>ETP<br>NA | | |
| KPC-2 | ✓ | ✓ | ✓ | ✓ |  |  |
| VIM-1 | ✓ | ✓ | ✓ | ✓ |  |  |
| SHV-200 | ✓ | ✓ | ✗ | ✓ |  |  |
| TEM-1 | ✓ | ✓ | ✓ | ✗ |  |  |

Cells in green color and marked with a ✓ indicates that the protein was identified with an $E$ -value $\leq 0.01$ , indicating high confidence in the identification. Cells in yellow color and marked with a ✓ indicates that the protein was identified with an $0.01 < E$ -value $\leq 1$ , indicating low confidence in the identification. Cells with ✗ indicates that the protein was not identified for that sample number (SN); cells with no mark indicates that the protein was not identified in that dataset; CTX, Cefotaxime; ETP, Ertapenem; MEM, Meropenem; NA, No Antibiotic. cases of no identification with no mark.

For bacterial cultures exposed to an antibiotic, one would expect the bacteria to express some of its AR proteins at high levels [96, 97]. MiCId’s workflow do identify, except for OXA-1, OXA-488 and OXA-905, all the predicted *β*-lactamase proteins. The AR protein OXA-1 is copresent with OXA-48, CTX-15 and TEM-1 in the genome of *K. pneumoniae* CCUG 70742; OXA-488 is present along with OXA-35, VIM-4, and PDC-35 in the genome of *P. aeruginosa* CCUG 51971; and OXA-905 is copresent with PDC-8 in the genome of *P. aeruginosa* CCUG 70744. For *K. pneumoniae* CCUG 70742, MiCId’s workflow identified OXA-48 in 3 samples via OXA-232, OXA-199 and OXA-548 with the 3 proteins being highly homologous to OXA-48 and having length normalized BLAST bit-scores of 2.04, 2.05 and 1.69 respectively. For *P. aeruginosa* CCUG 51971 MiCId’s workflow identified OXA-35 in 5 samples with high-confidence via OXA-19, OXA-101, OXA-35, OXA-147 and OXA-240 with the 5 proteins being highly homologous to OXA-35 and having length normalized BLAST bit-scores of 2.04, 2.03, 2.05, 2.03, and 1.98 respectively. The complete list of identified AR proteins can be found in Table S6.

MiCId’s identification results for *P. aeruginosa* CCUG 51971 and *P. aeruginosa* CCUG 70744 correlate well with a previous study showing that, in the model strain *P. aeruginosa*—PAO1 the gene of the OXA-50-like oxacillinase— is expressed at relatively low levels and is not inducible by *β*-lactams, while the gene of *bla*_PDC_, also expressed at relatively low levels usually, is strongly induced by *β*-lactams [98]. This could be the reason why MiCId did not detect OXA-488 in *P. aeruginosa* CCUG 51971 and OXA-905 in *P. aeruginosa* CCUG 70744 but detected PDC-35 in *P. aeruginosa* CCUG 51971, albeit only at the highest concentrations of meropenem. There are also several experimental reasons ranging from digestion enzyme, data dependent acquisition mode selection, protein expression level, as well as non-optimal liquid-chromatography separation that can be used to explain why some of the *β*-lactamase proteins were not identified or were not confidently identified. To further validate that the missed identification of *β*-lactamase proteins was not due to MICId’s inability, we analyzed all the 27 MS/MS experimental datafiles using the Proteome Discoverer software (version 2.4) and the results obtained, displayed in Table S9, are in agreement with MiCId’s results.

This assessment shows that MiCId’s workflow has a typical sensitivity value around 85% (and with a bound at about 72.7%), suggesting that it is a useful tool for the detection of AR proteins.

### 3.3 Using MiCId’s workflow to Investigate Possible Mechanisms of Antibiotic Resistance

We demonstrate here how MiCId’s workflow may help investigation of the possible mechanisms of antibiotic resistance of a human pathogen, *Pseudomonas aeruginosa* strain CLJ3, and compare the mechanism suggested by using MiCId with published results [72]. *Pseudomonas aeruginosa* strains were obtained from a patient having hemorrhagic pneumonia but treated unsuccessfully with antibiotics. Strain CLJ1, sensitive to antibiotics, was isolated before antibiotic therapy started; twelve days after antibiotic therapy started, as the patient conditions worsened, strain CLJ3 was isolated. Multiomics approach was used to understand the process of antibiotic resistance development in CLJ3. Genomics data shows that the genome of CLJ3, when compared to genome of CLJ1 has acquired several genetic modifications that could have contributed to phenotypic changes. Genomics data shows that antibiotic resistance of CLJ3 is probably linked to interruption-causing insertions detected in genes *oprD* and *ampD* [72].

For each strain, proteomics samples comprising of proteins contained in the whole-cell (W), inner and outer membranes (M), and secretome (S) were collected and used for MS/MS analysis [72]. The CARD database was used as the input AR protein database in MiCId’s workflow as it contains proteins belonging to multidrug efflux pumps [99]. The suggested mechanisms of antibiotic resistance for CLJ3 by using MiCId’s workflow agrees with the published results [72]. Comparing the AR proteins identified in membrane samples from CLJ3 and CLJ1, one notes that CLJ3 does not express the outer membrane protein *oprD* and is over-expressing the *β*-lactamase PDC. Lack of the outer membrane protein *oprD*, caused by interruption of *oprD* gene, makes the cell impermeable to most antibiotics in the *β*-lactam class [72,100,101]. Interruption in the *ampD* gene brings about the over-expression of *bla*_PDC_ as the *ampD* gene is responsible for the regulation of *bla*_PDC_ [102,103].

Table S10 contains the identifiers of AR protein families for each strain. Table S10 also shows that in agreement with the previous study on the mechanism of antibiotic resistance for CLJ3 was only obtained for the membrane samples. One obvious reason is that one expects to find higher concentration of *oprD* proteins in the membrane extract and of *β*-lactamase proteins in the periplasm, which is the cellular component between the inner- and outer-membrane of gram-negative bacteria, and thus can often be a component contaminant for the membrane samples [104,105]. This accentuates the necessity of sample extraction selection and sample fractionation when investigating possible mechanisms of antibiotic resistance [105–107].

We further used principal component analysis (PCA) to demonstrate the reproducibility of MiCId’s workflow. The vector component for each sample was set to be ln(1/*E*-value) of the identified AR protein family. AR proteins not identified in a given sample was assigned the *E*-value 100, yielding a vector component value of −4.605. For each sample, the components are further scaled to have norm 1. Figure 4 shows tight clusters for samples derived from whole-cell and membrane for each strain; for the secretome samples, data points are not as close, indicating that the secretome might not be suitable for studying AR proteins. The results from principal component analysis validates the reproducibility of MiCId’s workflow in AR protein identifications as shown by the tight sample clusters in Figure 4.

**Fig. 4:**
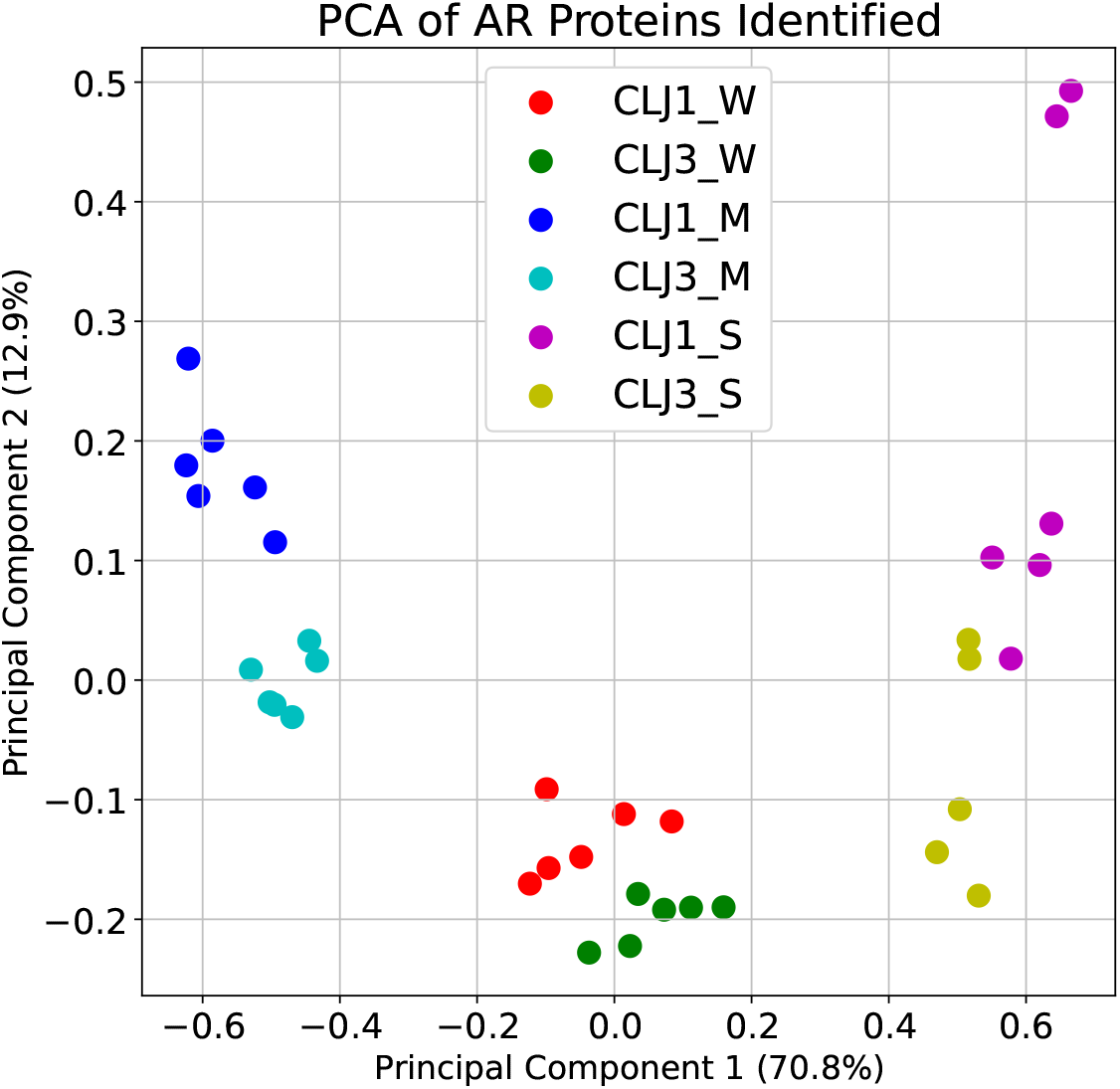
Principal component analysis (PCA) for antibiotic resistance (AR) protein families identified by MiCId’s workflow. Included in the PCA are 35 identification results, each from an experiment whose sample contains either *Pseudomonas aeruginosa* strain CLJ1 or *Pseudomonas aeruginosa* strain CLJ3 with proteins collected from whole-cell (W), membrane (M), or secretome (S). Also revealed in the plot, there are only five experimental replicates for the combination CLJ3-S, the other five combinations each has six experimental replicates. This brings the total number of experiments included to 35.

### 3.4 Execution Time of MiCId’s Workflow

With speed a main consideration, MiCId was written in C++ and its routines for organism and protein identifications were implemented using parallel programming. Hence, MiCId allows users to specify the desired number of cores for each job. Using 28,150 MS/MS spectra to query two databases of sizes 100GB (12,703 organisms) and 20GB (3,868 organisms), we measured the execution time of MiCId’s workflow in performing organism identification, biomass estimation, and protein identifications. Figure 5 shows that in the 20GB database takes about 13 min with 4 cores and reduces to around 6 min with 16 cores. On the other hand, the execution time in the 100GB database ranges from 17 min (with 4 cores) down to 7 min (with 16 cores). Our results indicate that when the database size increases by a factor of 5.0, the execution time increases only by a factor of about 1.2 (using 16 cores). This reflects the scalability of MiCId in handling large databases. Figure 5 also shows that the execution time reduction reaches a plateau at around 16 cores. This is because the C++ routine used to compute statistical significance for identified organisms and proteins is not yet parallelized, incurring a constant time cost. Table S2 contains the taxonomic identifiers for all the organisms in the 100GB and the 20GB databases as well as the identifiers for proteins taken from the ResFinder database.

**Fig. 5:**
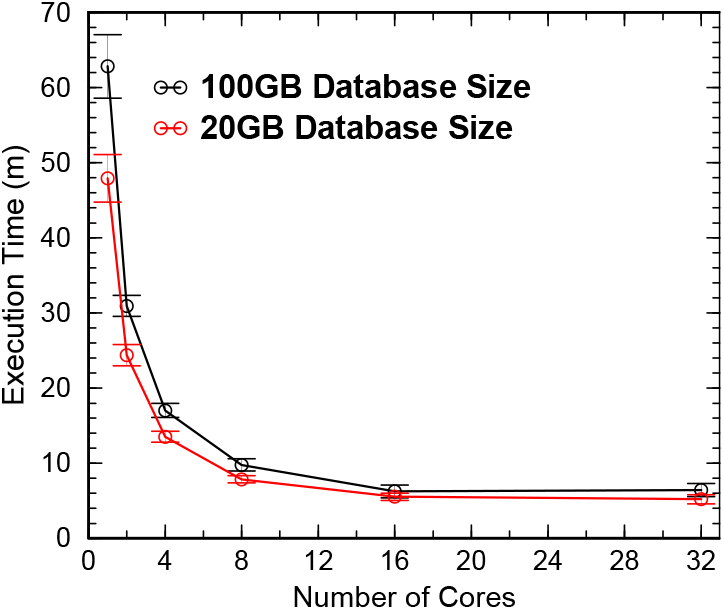
Average execution time, in minutes, of MiCId’s workflow in performing organism identification, biomass estimation, and protein identifications in a 100GB (containing 12,703 organisms) database and a 20GB (containing 3,868 organisms) database. There are 28,150 MS/MS spectra used as the queries. Results from using various number of cores are displayed. MiCId’s workflow execution time performance was carried-out in a computer running the operating system CentOS Linux release 7.9.2009 and containing 32 Intel(R) Xeon(R) central processing units (CPUs) with a clock speed of 2.60 GHz.

MiCId’s workflow execution time performance was done in a computer running the operating system CentOS Linux release 7.9.2009 and containing 32 Intel(R) Xeon(R) central processing units (CPUs) with a clock speed of 2.60 GHz. More information about the operating system and CPUs used is provided Table S11.

## 4 Conclusion

Fast and accurate identification of pathogenic bacteria along with the identification of AR proteins is of paramount importance for patient treatments and public health. The newly augmented MiCId workflow was designed to achieve this important goal by identifying AR proteins when processing MS/MS data acquired in high-resolution mass spectrometers. The augmented workflow of MiCId also fills the need for having mass spectrometry-based workflow for identifying bacteria along with AR proteins. We have shown in section 3.1 that the strategy employed by MiCId’s workflow for identifying AR protein yields sensible results. MiCId’s workflow identifies 93.5% (131/140) of the AR proteins that are also identified if the target protein database used is composed of protein sequences from the correct strain. Results from our AR protein identification assessment show that MiCId’s workflow has a sensitivity of 85% (with a lower bound at about 72.7%) and a precision of 95% when the *E*-value cutoff of 0.01 is used to control the number of false positives. Being fast, yielding sensible results, and having high sensitivity and high precision, MiCId is shown to be a valuable tool for identification of bacteria and their AR proteins.

The augmented workflow of MiCId is a self-contained tool capable of performing microorganism identification, protein identification, biomass estimation and AR protein identification in minutes using limited amount of computer resources available in most desktop and laptop computers. MiCid’s workflow was tested under (i) CentOS Linux release 7.9.2009, (ii) Red Hat Enterprise Linux Server release 7.9, (iii) Ubuntu release 18.04.3, and (iv) Windows 10 using Oracle VirtualBox 6.1.22 running Ubuntu release 18.04.3. Having a user-friendly graphical user interface, the new MiCId version (v.07.01.2021) for Linux environment is freely available for download at https://www.ncbi.nlm.nih.gov/CBBresearch/Yu/downloads.html.

## Acknowledgements

We thank the administrative group of the National Institutes of Health Biowulf Cluster, where all the computational tasks were carried out for the MiCId workflow. We thank the staff of the Culture Collection University of Gothenburg (CCUG, Gothenburg, Sweden) for providing bacterial strains. The CCUG is supported by the Department of Clinical Microbiology, Sahlgrenska University Hospital and the Sahlgrenska Academy of the University of Gothenburg, Sweden. RK, DJL, BPI, FSS and ERBM acknowledge support and funding from the Center for Antibiotic Resistance Research (CARe, Sahlgrenska Academy, University of Gothenburg). We thank the Proteomics Core Facility at the Sahlgrenska Academy, University of Gothenburg, for performing proteomics experiments and proteomics analysis using the Proteome Discoverer software version 2.4 (Thermo Fisher Scientific). This work was supported by the Intramural Research Program of the National Library of Medicine. Funding for Open Access publication charges for this article was provided by the National Institutes of Health.

## Electronic supplementary material

The online version of this article (….) contains supplementary material, which is available to authorized users.

**Supplementary File S1.** This Excel file contains relevant materials and results used in the manuscript. Figure S1 BLAST bit-score and length-normalized BLAST bit-score histograms. Figure S2 Length-normalized BLAST bit-score cutoff learning. Table S1 information about MS/MS files. Table S2 list of organisms and proteins used to build MiCId’s microorganismal databases and protein databases. Table S3 average similarity between *β*-lactamase protein families. Table S4 list of protein sequence identifiers for the correct bacteria strains and for the reference/representative strains. Table S5 list of antibiotic resistance proteins identified by MiCId’s workflow for sample number 1-62 when using as database proteins from the correct strain plus proteins from the ResFinder database, and proteins from the reference/representative strain plus proteins from the ResFinder database. Table S6 list of antibiotic resistance proteins identified by MiCId’s workflow for sample number 1-126 when using as database proteins from the reference/representative strain plus proteins from the ResFinder database. Table S7 species level identification for sample number 1-126. Table S8 *β*-lactamase proteins and their target *β*-lactam drug classes for the six strains cultivated with *β*-lactam used in our study. Table S9 list of antibiotic resistance proteins identified by Proteome Discoverer software version 2.4 (Thermo Fisher Scientific) for sample number 1-27. Table S10 list of antibiotic resistance proteins identified by MiCId;s workflow for sample number 28-62 when using as database proteins from the reference/representative strain plus proteins from the CARD database. Table S11 provides information about the computer operating system and CPUs used to measure MiCId’s workflow execution time.

